# Sensorimotor awareness requires intention: Evidence from minuscule eye movements

**DOI:** 10.1101/2024.07.02.601661

**Authors:** Jan-Nikolas Klanke, Sven Ohl, Martin Rolfs

## Abstract

Microsaccades are tiny eye movements that are thought to occur spontaneously and without awareness but can also be intentionally controlled with high precision. We used these tiny visual actions to investigate how intention modulates sensorimotor awareness by directly comparing intended, unintended, and spontaneous microsaccades. In addition, we dissociated the effects of action intention and the actions’ visual consequences on awareness. In 80% of all trials, we presented a stimulus at high temporal frequency rendering it invisible during stable fixation. Critically, the stimulus became visible when a microsaccade in the same direction caused it to slow down on the retina (generated microsaccade condition; 40% of trials) or when the microsaccades’ visual consequence was replayed (replayed microsaccade condition; 40% of trials). Participants reported whether they perceived the stimulus (visual sensitivity), whether they believed they had made a microsaccade (microsaccade sensitivity), and their level of confidence that their eye movement behavior was linked to their perception (causality assignment). Visual sensitivity was high for both, generated and replayed microsaccades and comparable for intended, unintended, and spontaneous eye movements. Microsaccade sensitivity, however, was low for spontaneous microsaccades, but heightened for both intended and unintended eye movements, showing that the intention to saccade or fixate enhances awareness of otherwise undetected eye movements. Visual consequences failed to aid eye movement awareness, and confidence ratings revealed a poor understanding of a causal relationship between eye movement and sensory consequence. These findings highlight the functional relevance of intention in sensorimotor awareness at the smallest scale of visual actions.

**Significance statement:** While eye movements are among the most frequent human actions, they are rarely perceived consciously, despite causing sweeping changes in retinal inputs. Here investigate how intention can modulate awareness of even the smallest human actions: microsaccades. We developed a novel paradigm that allowed us to dissociate the role of action intention and an action’s sensory consequence for awareness, two factors that previous research has typically confounded. Our data provide strong evidence that observers can detect small eye movements reliably and demonstrates that sensitivity towards microsaccades was neither driven by an eye movement’s motor component nor its visual consequences alone. Instead, we find that intention opens a gate to sensorimotor awareness, even for actions typically too small to be perceived.

## Introduction

Vision is inherently active (Ahissar & Arieli, 2001; Rolfs, 2015; Rucci et al., 2018; Rucci & Victor, 2015)—the eyes move incessantly to sample different aspects of the environment over time. Despite the high frequency of these visual actions and their immediate visual consequences (Rolfs & Schweitzer, 2022), we appear to have little access to our own past or ongoing eye movement behavior (Marti et al., 2015; Võ et al., 2016). It thus remains elusive to what degree we have sensorimotor awareness, or even a sense of agency (Haggard, 2017), for eye movements at all. Sensorimotor awareness likely hinges on the degree of intended control over these movements and the distinction between self-generated and externally-generated sensory signals. But these two factors are inherently difficult to manipulate in any domain of action control, as one must exactly match intended and unintended movements with respect to both their kinematics and their sensory consequences. Here, we address this challenge by capitalizing on microsaccades—minuscule eye movements with reliable kinematics that occur spontaneously during gaze fixation (Cook et al., 1966; Yarbus, 1967; Zuber et al., 1965), but can also be controlled (Guzhang et al., 2024; Ko et al., 2010; Poletti et al., 2020; Shelchkova & Poletti, 2020; Willeke et al., 2019)—to investigate how (1) the intention to move and (2) the resulting visual consequences modulate sensorimotor awareness for eye movements.

Microsaccades are an intriguing oculomotor model for eye movement awareness that allows us to disentangle these two factors. First, microsaccades frequently occur spontaneously when observers have the intention to fixate. Given their miniscule size, they are assumed to escape awareness (Engbert & Kliegl, 2004; Martinez-Conde et al., 2004; Rolfs, 2009; Rosenzweig & Bonneh, 2019). At the same time, observers can intendedly move their eyes over similarly small amplitudes when guided by visual cues (Ko et al., 2010; Poletti et al., 2020) or memory alone (Hafed & Goffart, 2020; Willeke et al., 2019). This provides an experimental handle on the effect of intention on eye movement awareness: By directly comparing awareness for spontaneous, unintended, and intended microsaccades, we can assess if the intention to move makes observers more sensitive to self-generated actions. Second, microsaccades lead to small, rapid displacements of the visual scene on the retina that are not perceived under normal viewing conditions. Fast flickering (Deubel & Elsner, 1986) or phase-shifting stimuli (Kelly, 1990), however, render minute eye movements visible by painting their immediate sensory consequences on the retina. This allows us to carefully manipulate the presence and magnitude of the visual consequence of the eye movement and uncover their impact on sensorimotor awareness.

So far, only one published report investigated subjective awareness of microsaccades (Haddad & Steinman, 1973), and they were never compared directly. Haddad and Steinman (1973) discovered that expert observers can detect spontaneous microsaccades but fail to recognize their direction. However, it remained unclear if microsaccades were ever falsely reported in that study. The rate of false alarms, however, is required to determine observers’ sensitivity (Green & Swets, 1966).

We developed a paradigm that directly addresses if and how eye movement awareness depends on the observer’s intention to move and the resulting retinal consequence. To investigate movement intention, we directly compared observers’ sensitivity towards having generated intended, unintended (**Experiment 1**), or spontaneous microsaccades (**Experiment 2**). In **Experiment 1**, we instructed observers to either execute a small, deliberate saccade to a memorized target location (Willeke et al., 2019) as soon as the fixation point (and saccade targets) disappeared (instructed saccade trials; **Fig. 1a**), or to maintain fixation (instructed fixation trials; **Fig. 1a**). Saccades executed in saccade trials were labelled *intended microsaccades*. Conversely, saccades executed in fixation trials were labelled *unintended microsaccades*. In our second experiment, observers were informed about the existence and visual consequences of microsaccades in our paradigm but did not receive specific instructions regarding a required eye movement behavior (**Fig. 1a**). Thus, we labelled any occurring saccades as *spontaneous microsaccades*.

**Figure 1.**
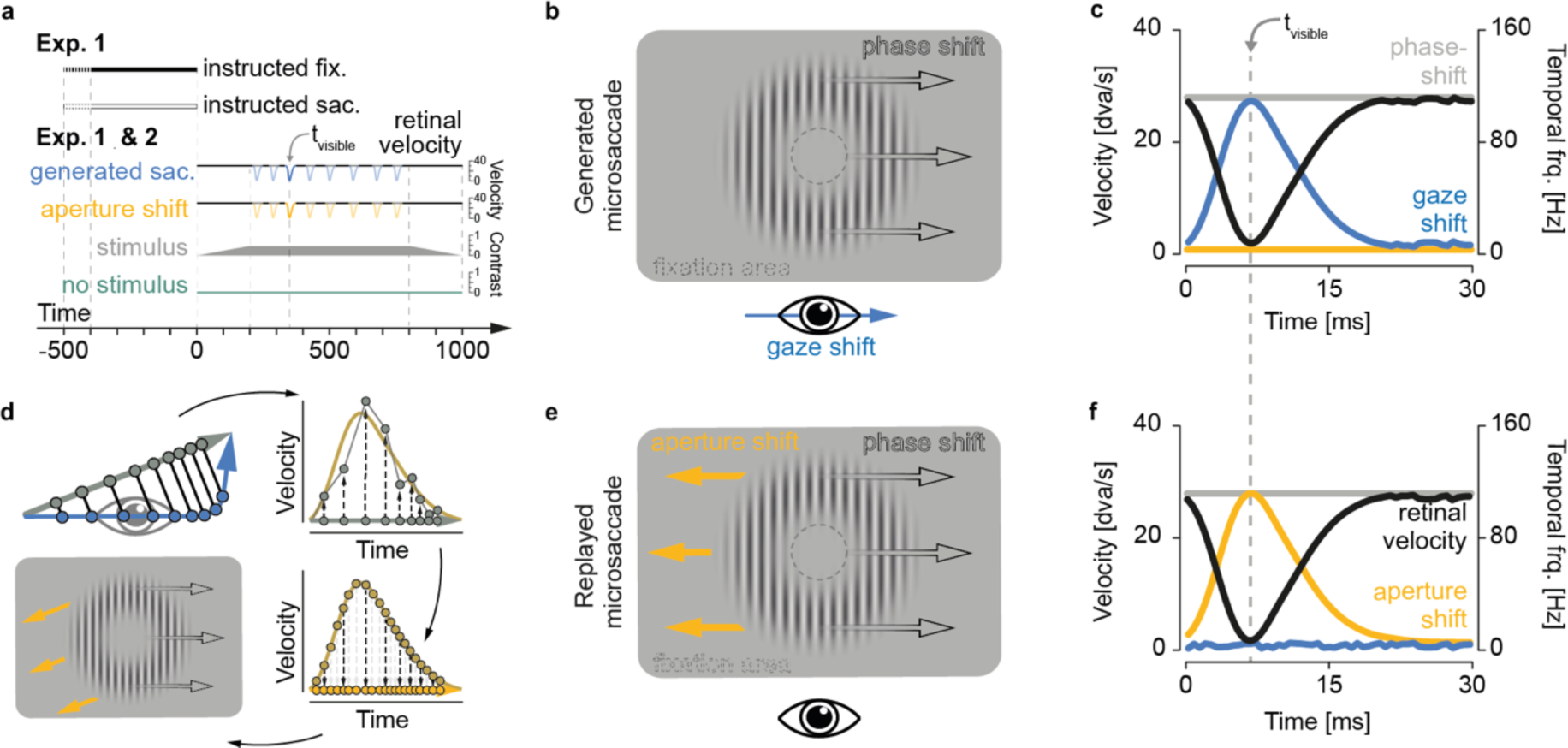
Experimental protocol and stimulus design. **a** Procedure in Experiments 1 and 2. Bars (Experiment 1) indicate presentation of the fixation dot and saccade target in trials in which either an intended (white) or an unintended microsaccade (black) was prompted. Black lines indicate constant retinal velocity of the stimulus, colored sections denote the stimulus being slowed down on the retina by a generated (blue) or replayed (yellow) microsaccades. Trapezoid shape depicts contrast modulation of the stimulus (grey). We included a stimulus absent condition (green) as an additional control, in which the stimulus contrast was set to zero. **b** Stimulus display for generated microsaccades. Gray arrows indicate the direction of the phase shift, blue arrow indicates the direction of a microsaccade that leads to a retinal stabilization of the stimulus. **c** Velocity profiles of the phase shift (gray line), gaze shift (blue line), aperture shift (yellow line), and retinal velocity (black line) for generated microsaccades. The phase shift leads to temporal frequencies >60 Hz and renders the stimulus invisible during fixation. Only if the stimulus is slowed down on the retina by a microsaccade, will the combined stimulus velocity drop below the detection threshold. **d** Schematic depiction of the steps to generate the aperture motion that replays the retinal consequence of a previous microsaccade (clockwise): Projection of sampled gaze position to saccade vector (upper left), fitting of a gamma function to the velocity profile along the saccade vector (upper right), recalculation of the gaze positions along the saccade vector based on the fitted velocities (lower right), aperture shift in the opposite direction to mirror retinal image displacement (lower left). **e** Stimulus display for replayed microsaccades. Gray arrows indicate the direction of the phase shift (same as in b), yellow arrows indicate the direction of an aperture shift that replays the retinal consequence of a microsaccade and leads to a comparable retinal stabilization of the stimulus. **f** Velocity profiles of replayed microsaccades. Colors are same as in c. If the stimulus is slowed down on the retina by a replayed microsaccade (i.e., the aperture shift), will the combined stimulus velocity drop below the detection threshold.

To investigate the role of visual consequence on eye movement awareness, we presented a high-temporal frequency stimulus that was invisible during fixation (> 60 Hz, cf. Castet & Masson, 2000), but rendered visible when microsaccades with matching kinematics briefly stabilized it on the retina (cf. Deubel et al., 1987; Deubel & Elsner, 1986; Kelly, 1990; **Fig. 1b/c**). We added a condition in which the stimulus’ aperture replayed a previous eye movement back to the observer (**Fig. 1d/e/f**), such that the observer could not determine the presence of a microsaccade just based on the visual information alone. Finally, we included a condition in which stimulus’ contrast was set to 0 to compare detection of eye movements that did not cause any visual consequence to eye movements that did. For each of observer, we determined three types of sensitivity: (1) their visual sensitivity for detecting a brief visual stimulus contingent on microsaccades or their replayed sensory consequences (visual sensitivity), (2) their ability to report whether they generated a microsaccade (microsaccade sensitivity) and its contingency on stimulus presence, and (3) their confidence that their eye movement behavior was linked to their perception (causality assignment).

## Results

### Motor control for microsaccades

The rate of unintended microsaccades was significantly lower than the rate of intended microsaccades (*t* (9) = 3.49, *p* = 0.007; unintended: mean = 0.11 s^-1^±0.09; intended: mean = 0.38 s^-1^±0.21; **Fig. 2a**), confirming that participants can, to some degree, control microsaccadic behavior. Target distance (ranging from 0.2 dva to 1 dva) affected the ability to generate intended microsaccades (one-way rmANOVA: *F* (4,36) = 4.84, *p* = 0.003; **Fig. 2a**), with increasing microsaccade rates for larger target distances (0.2 dva: 0.22 s^-1^±0.15; 0.4 dva: 0.38 s^-1^±0.23; 0.6 dva: 0.43 s^-1^±0.25; 0.8 dva: 0.44 s^-1^±0.24; 1.0 dva: 0.4 s^-1^1±0.22). The rate for spontaneous microsaccades from **Experiment 2** (0.19 s^-1^±0.13; **Fig. 2a**) was in between those of unintended and intended microsaccades, and not statistically different from either intended (*t* (15.2) = 1.76*, p* = 0.098), or unintended (*t* (16.4) = –1.09, *p >* 0.250) ones (see section *Saccade rates* in **Supplementary material**). Also, amplitudes of intended microsaccades from **Experiment 1** increased monotonically (0.2 dva: 0.41±0.08 dva; 0.4 dva: 0.50±0.10 dva; 0.6 dva: 0.56±0.10 dva; 0.8 dva: 0.64±0.10 dva; 1 dva: 0.66±0.11 dva); **Fig. 2b**) with target distance (*F* (4,36) = 28.40, *p* < 0.001). A linear mixed effects model, fit to the amplitudes of those intended eye movements, revealed significant positive estimates for all successive difference contrasts (all *p*s < 0.001, except 0.8 vs. 1.0 dva: *t* (3238.6) = 1.75, *p* = 0.079, beta = 0.02±0.02). This result suggests that observers adapted their microsaccade amplitudes to the target distances (for more information see section *Accuracy and precision of intended microsaccades* in the **Supplementary material**).

**Figure 2.**
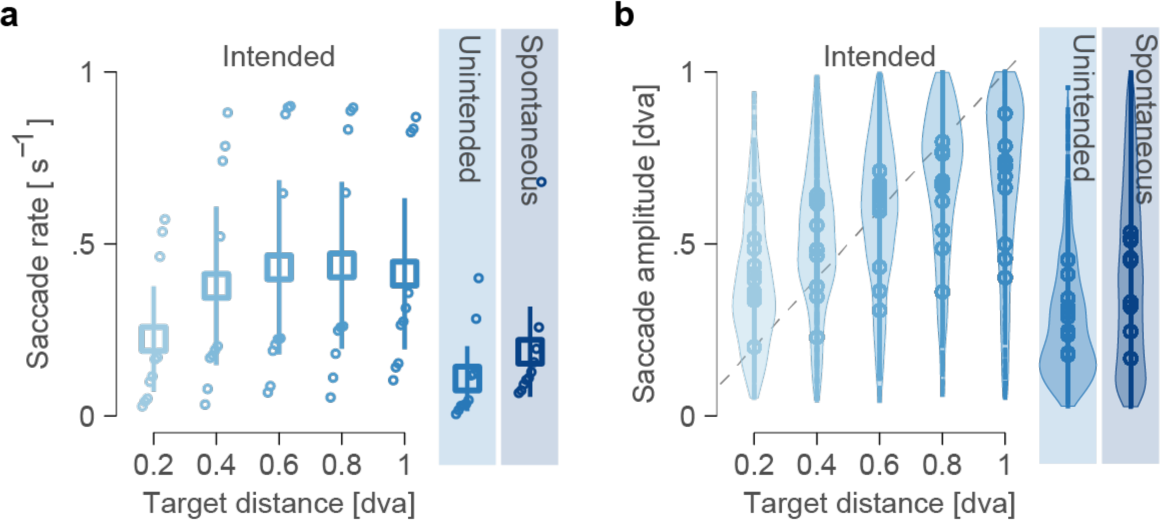
Accurate control of microsaccades. **a** Rates of different types of microsaccades and, for intended microsaccades, different target distances. Empty circles denote average rates per participant and condition, sorted from lowest to highest by value, squares indicate group means. Error bars show 95% confidence intervals. **b** Amplitudes of different types of microsaccades and, for intended microsaccades, different target distances (0.2*–*1 dva). Empty dots indicate average amplitudes per participant and target amplitude, violin-plots indicate distribution of all saccade amplitudes and target distances.

### Visual sensitivity to intra-saccadic stimulation

Next, we confirmed that observers were indeed visually insensitive to the high-temporal frequency stimulus displayed during fixation: In the absence of microsaccades, observers’ sensitivity for detecting the stimulus was not statistically different from 0 (**Exp. 1:** d’ = 0.16±0.19; **Exp. 2:** d’ = 0.10±0.29; **Fig. 3a**). This insensitivity was indistinguishable between intended (d’ = 0.13±0.18) and unintended (d’ = 0.20±0.22; **Exp. 1**: *t* (9) = –1.13, *p* > 0.250) as well as spontaneous microsaccades compared to eye movements from **Experiment 1** (**Exp. 1 vs 2:** *t* (15.4) = 0.44, *p* > 0.250).

**Figure 3.**
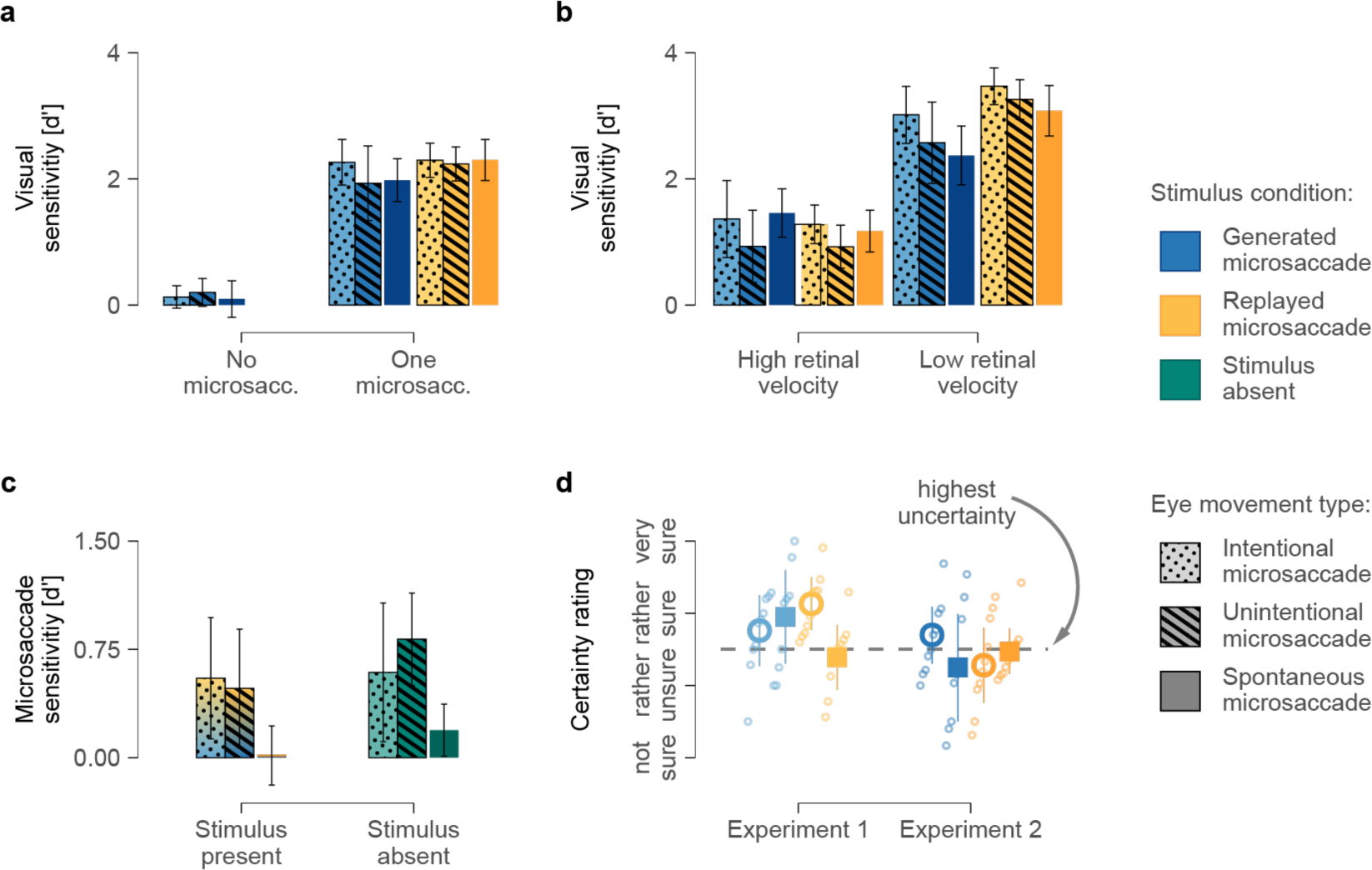
Visual and microsaccade sensitivity. **a** Visual sensitivity as a function of microsaccade generation for different stimulus conditions and eye movement types. **b** Visual sensitivity as a function of retinal velocity (in low-velocity trials, the saccade’s peak velocity was within ±30 dva/s of the grating’s velocity; in high-velocity trials, it was outside that range). **c** Microsaccade sensitivity as a function of stimulus presence and eye-movement type. **d** Certainty of judgment about the causal relationship between stimulus percept and eye movement for the two experiments. Error bars indicate 95% confidence intervals.

Intended and unintended microsaccades (d’ = 2.10±0.46) as well as their replayed retinal consequences (d’ = 2.27±0.25), rendered the stimulus highly visible (**Fig. 3a**). A two-way rmANOVA revealed that the increase in visibility was the same irrespective of stimulus condition (generated vs. replayed; *F* (1,9) = 1.61, *p* = 0.236) and eye-movement type (intended vs. unintended; *F* (1,9) = 4.15, *p* = 0.072; interaction: *F* (1,9) = 2.34, *p* = 0.160). In **Experiment 2**, spontaneous microsaccades, both generated (d’ = 1.98±0.34) or replayed (d’ = 2.30±0.33), resulted in a similar visual sensitivity (**Fig. 3a**). A mixed-measures ANOVA with the between-subject factor experiment (**Exp. 1** vs. **2**) and the within-subject factor stimulus condition (generated vs. replayed) revealed that stimulus sensitivity was not significantly different between the two experiments (*F* (1,18) = 0.04, *p* > 0.250). A significant main effect of stimulus condition (*F* (1,18) = 8.87, *p* = 0.008) in the absence of an interaction (*F* (1,18) = 0.84, *p* > 0.250), however, showed that stimulus sensitivity was slightly higher when the sensory consequences of microsaccades were replayed (d’ = 2.22±0.20) rather than generated by an eye movement (d’ = 1.98±0.28).

The match between microsaccade kinematics and stimulus parameters markedly affected visual sensitivities (**Fig. 3b**). Intended and unintended microsaccades from **Experiment 1** that matched the speed and direction of the high-temporal frequency stimulus— leading to low retinal velocities within ±30 dva/s of the grating’s velocity—yielded significantly higher visual sensitivity (d’ = 3.08±0.34) than trials with mismatching parameters (d’ = 1.12±0.35; **Exp. 1:** *t* (8) = –13.94, *p* < 0.001). A three-way rmANOVA revealed that both retinal velocity (low vs. high velocity; *F* (1,8) = 194.39, *p* < 0.001) and stimulus condition (generated vs. replayed; *F* (1,8) = 5.94, *p* = 0.041) affected visibility; sensitivity was higher for low compared to high retinal velocities, and for replayed compared to generated microsaccades. A significant interaction between retinal velocity and stimulus condition (*F* (1,8) = 7.08, *p* = 0.029) highlighted that the impact of retinal velocity on visibility was larger for replayed than generated microsaccades. Lastly, a significant main effect of eye movement type (*F* (1,8) = 5.87, *p* = 0.042) revealed that stimulus visibility was also slightly higher for intended (d’ = 2.28±0.30) compared to unintended microsaccades (d’ = 1.92±0.39)—potentially due to the overall lower size and peak velocity of unintended microsaccades (see section *Parameters of different eye movement types* in **Supplementary material**). All remaining interactions were not significant (all *p*s > 0.250).

Spontaneous microsaccades in **Experiment 2** showed a similar benefit in visual sensitivity for low (d’ = 2.73±0.41) over high (d’ = 1.32±0.33) retinal velocities (**Exp. 2:** *t* (9) = –8.28, *p* < 0.001; **Fig. 3b**). In a three-way mixed-measure ANOVA with experiment as the between-subject factor, and stimulus condition as well as retinal velocity as within-subject factors, the main effect of retinal velocity was highly significant (*F* (1,17) = 226.33, *p* < 0.001) while the main effect of experiment was not (*F* (1,17) = 0.17, *p* > 0.250). Thus, the increase in visual sensitivity for low compared to high retinal velocities was consistent for all three types of eye movements. Similarly, stimulus condition (generated vs. replayed) affected visual sensitivity (*F* (1,17) = 11.93, *p* = 0.003) and interacted with retinal velocity (*F* (1,17) = 33.07, *p* < 0.001), implying that the general advantage for low compared to high retinal velocities is larger for the replayed than the generated stimulus condition. Finally, a significant interaction between the experiment and the retinal velocity (*F* (1,17) = 5.93, *p* = 0.026) indicated that the gain in visibility for low compared to high retinal velocities is smaller for spontaneous than for intended and unintended microsaccades. No other interactions were significant (all *p*s > 0.191).

The analysis of visual sensitivity confirmed that our high-temporal frequency stimulus was visible only during the presence of generated or replayed microsaccades, with only small variations between instruction and, thus, eye-movement types. Sensitivity increased when stimulus and microsaccade parameters matched, confirming that gaze-contingent retinal stabilization determined the visibility during the microsaccade. Additionally, observers were slightly more sensitive towards replayed compared to generated microsaccades. We attribute this difference to an overestimation of saccade peak velocity in video-based eye tracking (cf. Schweitzer & Rolfs, 2022), which would yield a small but systematic discrepancy in the effective retinal velocity between the generated and replayed condition. In addition, saccadic suppression (i.e., the decrease in visual sensitivity during eye movements) may have reduced visual sensitivity as well (Hafed & Krauzlis, 2010; Zuber & Stark, 1966).

### Microsaccade sensitivity

We next examined how sensitive observers were in detecting their own eye movements and how this microsaccade sensitivity depended on the absence vs. presence of a visual stimulus. In **Experiment 1**, observers were moderately sensitive to both intended (d’ = 0.57±0.43), and unintended microsaccades (d’ = 0.65±0.33; **Fig. 3c**). A two-way rmANOVA revealed that sensitivity was comparable between the two different types of eye movements (intended vs. unintended; *F* (1,9) = 0.11, *p* > 0.250). The presence of a visual stimulus significantly decreased microsaccade sensitivity (present vs. absent; *F* (1,9) = 5.40, *p* = 0.045). Observers were more sensitive to their own microsaccades in trials in which no stimulus was present (d’ = 0.71±0.28) as compared to trials with a stimulus (d’ = 0.51±0.30; **Fig. 3c**). The interaction of eye-movement type and stimulus presence, on the other hand, was not significant (*F* (1,9) = 2.64, *p* = 0.139).

Next, we compared the results for intended and unintended microsaccades to spontaneous eye movements from **Experiment 2**. In line with our predictions, we found microsaccade sensitivity to be much lower for spontaneous microsaccades both in stimulus absent (d’ = 0.19±0.18) and stimulus present trials (d’ = 0.02±0.21; **Fig. 3c**). Indeed, in stimulus present trials, microsaccade sensitivity for spontaneous microsaccades was indistinguishable from 0. A two-way mixed-measures ANOVA that assessed microsaccade sensitivity based on stimulus presentation as a within-subject factor, and experiment as a between-subject factor revealed main effects of stimulus presentation (absent vs. present; *F* (1,18) = 8.31, p = 0.010) and experiment (*F* (1,18) = 12.91, p = 0.002), with no interaction (*F* (1,18) = 0.02, p > 0.250). Thus, stimulus-absent trials led to slightly higher microsaccade sensitivity in both experiments, and microsaccade sensitivity was lower for spontaneous compared to intended and unintended microsaccades.

In summary, observers were just as sensitive to unintended microsaccades during instructed fixation as to intended microsaccades following an instruction to move the eyes. In contrast, spontaneous microsaccades typically escaped awareness. Indeed, the subjective microsaccade-contingent change in stimulus visibility did not enhance microsaccade sensitivity (see section *Microsaccade sensitivity as a function of stimulus perception* in **Supplementary Material**). Our data instead support the opposite conclusion: the presence of a visual stimulus had a detrimental effect on microsaccade sensitivity.

### Causal assignment: Relating eye movements to their consequences

We investigated whether observers were able to detect if their own eye movements caused the stimulus to become visible. We predicted that, if microsaccades were made intendedly and consciously (and if observers understood that the stimulus became visible because of the microsaccade), observers should be confident that their eye movements caused the brief change in stimulus visibility. In other words, certainty about the causal link between eye movement and stimulus visibility should be a function of sensorimotor awareness of the eye movement and, thus, higher when the generation of an eye movement was correctly detected. Due to the lack of trials in which observers correctly detected unintended microsaccades that rendered the stimulus visible, we collapsed data for intended and unintended eye movements.

Focusing on **Experiment 1** first, we found that the reported levels of certainty were close to the scale’s mid-point (i.e., the highest level of uncertainty, 0; **Fig. 3d**) irrespective of whether the microsaccade was generated (0.30±0.41) or replayed (0.22±0.32). Observers did, however, show a slightly higher certainty for correct causality assignments (0.39±0.31) compared to when causality was assigned incorrectly (0.13±0.40; **Fig. 3d**), and a two-way rmANOVA confirmed the significance of this difference (*F* (1,8) = 8.96, *p* = 0.017). While the difference between the two stimulus conditions remained insignificant (generated vs replayed: *F* (1,8) = 0.47, *p* > 0.250), a significant interaction (*F* (1,8) = 5. 37, *p* = 0.049) indicated a higher certainty for correct (over incorrect) assignments only for replayed eye movements (0.72±0.36), not for generated ones (–0.20±0.61).

In **Experiment 2**, we observed comparable albeit slightly lower levels of certainty for generated (–0.05±0.42) and replayed spontaneous microsaccades (–0.09±34; **Fig. 3d**). Unlike for **Experiment 1**, observers were not more confident in assigning causality correctly, compared to incorrectly (*t* (8) = 2.19, *p* = 0.060; correct: 0.03±0.34; incorrect: –0.17±0.43; **Fig. 3d**), and a two-way rmANOVA showed neither factor nor their interaction to be significant (all *p*s > 0.06).

In summary, observers’ reports suggested a very limited understanding of the relationship between eye movement and the stimulus on a trial-by-trial bases. Notably, we observed an increase in certainty when participants were presented with replayed eye movements from the first experiment. This increase in certainty implies that observers were more confident about the absence than the presence of a link between the stimulus and the eye movement—and only when they were sensitive to their eye movements at all.

## Discussion

### The role of intention for sensorimotor awareness

We investigated how action intention and an action’s visual consequence affect sensorimotor awareness in human observers. We revealed that action intention enhances sensorimotor awareness even for movements that are typically too small to be perceived: Observers were significantly more sensitive to their microsaccades when they intended to make or avoid them compared to when such microsaccades occurred spontaneously (**Fig. 3c**). Our findings demonstrate that microsaccades, while phenomenally thin (Clark et al., 2013; Haggard, 2017) and prone to escape awareness when generated spontaneously (i.e., in the absence of an intention), can be recognized in principle, and at a level comparable to intended microsaccades. Importantly, an action’s sensory consequence did not lead to a similar increase in saccade sensitivity, therefore pointing towards action intention as the main factor for sensorimotor awareness.

In our study, we examined the role of movement intention by presenting instructions in the beginning of each trial, prompting observers’ intentions to either generate or suppress a microsaccade (**Exp. 1**). In a second experiment, we repeated the procedure but without providing explicit instructions to the observers (**Exp. 2**). The difference in sensitivity between intended (**Exp. 1**) and spontaneous microsaccades (**Exp. 2**) clearly demonstrates an effect of intention. Interestingly, the parameters of spontaneous and unintended microsaccades were similar in our experiments (see **Fig S2**), and the degree of sensorimotor awareness is not a function of movement parameters alone (e.g., amplitude). Taken together, our data caution against the classification of saccadic eye movements according to a system of distinct types based on fixed parameters (e.g., amplitude, duration, or latency) or levels of conscious processing: Our observers were sensitive to minuscule eye movements—irrespective of whether they were planned (like intended microsaccades, **Exp. 1**) or unplanned (like the unintended microsaccades, **Exp. 1**). However, in the absence of an intention, saccades of similar size, peak velocity, duration, and latency (i.e., spontaneous microsaccades from **Exp. 2**) routinely escaped conscious detection. Instead of a rigid typology of saccadic activity, our data support the idea of an oculomotor continuum along which saccades are generated (Hafed, 2011; Martinez-Conde et al., 2013; Rolfs et al., 2008; Zuber et al., 1965). Sensorimotor awareness of miniscule motor acts is, in line with this view, not pre-determined by the type of motor act, but by additional factors—most evidently, action intention.

### Motor control for minuscule eye movements

Generating instructed microsaccades in the absence of visual cues becomes more difficult the smaller the required amplitude is: We found lower saccade rates and reduced accuracy for smaller microsaccades which indicated that observers frequently overshot particularly small target distances (i.e., 0.2 and 0.4 dva; **Fig. 2a** and **Fig. S1a**). Despite that challenge, our data demonstrate that observers can generate small eye movements reliably—even in the absence of a foveated visual anchor: Microsaccades were more likely following an instruction to make a microsaccade and microsaccade amplitudes scaled with target distance. Successful execution of intended microsaccades increased with target distance, suggesting a graded control over minute eye movements as a function of saccade amplitude (cf. Willeke et al., 2019, 2022; see **Fig. 2b**). For trials in which observers were instructed to fixate, we revealed fewer and smaller microsaccades indicating that these eye movements were generated despite the intention to fixate. Their average latency also more closely resembled that of spontaneous microsaccades (**Fig. S2d**), indicating that unintended saccades are not a type of goal-directed saccade but rather saccadic intrusions (cf. Abadi & Gowen, 2004). Taken together, these results indicate that our observers exerted a high level of conscious control over their eye movement generation. But this control is not perfect: The small number of unintended microsaccades (**Fig. 2a**) suggests that some level of involuntary eye movement activity cannot be controlled—even when participants are explicitly instructed to do so.

The data provided by our two experiments is partially in line with previous findings claiming that expert observers can detect spontaneous microsaccades (Haddad & Steinman, 1973). Observers in our study showed no sensitivity towards spontaneous microsaccades but exhibited an increased sensitivity towards their saccades of the same size occurring when instructed to fixate (**Fig. 3c**). Assuming the expert observers in Haddad’s and Steinman’s original study received a similarly explicit instruction to fixate, we can assume a similarly heightened sensitivity towards unintended small eye movements as exhibited by our participants. Nevertheless, we want to offer an alternative interpretation of our respective results, which would accommodate that Haddad and Steinman genuinely measured the detection of spontaneous microsaccades. We find that detection (i.e., hit rates) of spontaneous microsaccades is significantly different from zero, when collapsing over all stimulus conditions and stimulus perception. At the same time, our observers exhibited a comparable increase in false alarm rates for spontaneous microsaccades (again irrespective of stimulus condition or perception; see section *Microsaccade (mis-) detection based on stimulus condition and perceptual report* in the **Supplementary material** for the extended analysis of hits and false alarms), rendering observer sensitivity (d’) towards spontaneous microsaccades not significantly different from zero (**Fig. 3c**). By focusing on hit rates only, Haddad and Steinman may have inadvertently misconstrued their reports that minuscule eye movements were generated (when indeed they were) as sensitivity towards microsaccades. Their observer’s inability to report the direction of the generated microsaccades can be seen as support for this interpretation of their data.

### Role of an action’s visual consequences for sensorimotor awareness

To investigate how an action’s visual consequence affects observers’ awareness of the underlying eye movement, we determined microsaccade sensitivity as a function of stimulus presence vs. absence. Interestingly, we found that observers were slightly more sensitive to their eye movements in trials in which the stimulus was absent rather than present (see **Fig. 3b**), suggesting that vision may have a detrimental effect on eye movement awareness. To examine this result more closely and explain a seemingly complex set of data, we directly compared hit and false alarm rates for microsaccades depending on eye movement type, stimulus condition, and perceptual reports (see section *Microsaccade (mis-) detection based on stimulus condition and perceptual report* in the **Supplementary material**).

We found higher hit rates in trials in which participants reported having perceived the stimulus compared to trials in which participants reported not having perceived it, suggesting that detection of small eye movements is heightened following a change in the visual input. Similar detection rates for replayed and generated microsaccades on the other hand suggest that a match between the visual consequence and the eye movement does not have to be perfect for observers to conclude that an eye movement has occurred. While this ostensibly counters our initial impression and instead suggests eye movement awareness benefits from the display of visual consequences, turning to false alarms levels this impression: We found that observers reported the erroneous belief to have generated a microsaccade significantly more often when a replayed eye movement was perceived by the observer compared to when it was not (**Fig. S4**). The increase in false alarms was comparable to the increasing hit rates and a re-analysis of microsaccade sensitivity based on stimulus perception (rather than presentation) revealed no significant difference between trials in which observers reported having perceived the stimulus compared to trials in which observers reported having perceived no stimulus (**Fig. S3**). Our findings thus demonstrate that while visual events strongly affect an observers’ beliefs about their eye movements, their effect on eye movement awareness are surprisingly limited.

However, this limited effect on awareness may well be an effect of our paradigm: To reveal how eye movement awareness was affected by intention, our paradigm decouples the presence of an eye movement and its visual consequences, as eye movements were neither necessary (cf. replayed microsaccades) nor sufficient (cf. no-stimulus condition) for seeing the stimulus. Seeing the stimulus, in turn, bore equally little information about eye movement generation: The stimulus was rendered visible in the absence of an eye movement when a visual consequence was replayed and the stimulus remained invisible irrespective of microsaccade generation in stimulus-absent trials. Relying on stimulus perception was, therefore, a poor strategy to try and gauge eye movement generation in the context of our paradigm. In everyday life, however, observers experience their eye movements predominantly as (highly predictable) changes in what we look at, arguably a visual change. The over-reliance of our participants on stimulus perception to estimate saccade generation (as evidenced by the high number of false alarms following stimulus perception for all but unintended microsaccades; cf. **Fig. S4**) indicates that our beliefs about eye movement generation critically relies on vision.

Lastly, why do we find a higher microsaccade sensitivity in stimulus absent compared to stimulus present trials? We argue that this is a combined effect of the observer’s tendency to over-estimate eye movement generation when they perceived the stimulus (**Fig. S4**) and the slightly higher stimulus sensitivity in replay condition trials (**Fig. 3a/b**). A higher visibility of the stimulus led to marginally higher false alarm rates that—together with comparable hit rates between stimulus conditions—led to a slightly lower sensitivity in stimulus present trials (**Fig. 3c**).

Taken together, our data support that visual consequences of eye movements are relevant for sensorimotor awareness of microsaccades: In a situation in which a minuscule eye movement is itself not very salient, human observers tend to use visual information to try and estimate if an eye movement was generated. While using vision would be a sound strategy under natural viewing conditions, where the immediate visual consequences of an eye movement are rarely matched by external visual events, our paradigm limited the utility of this approach: By adding a condition in which visual consequences of a saccade was replayed back to the observer, we decoupled eye movements and their visual consequences to reveal, for the first time, that movement intention is an important driver of sensorimotor awareness for minuscule eye movements.

### Causal assignment

Finally, we investigated if participants could relate their eye movements to stimulus perception after experimentally controlling for an action’s visual consequence. More specifically, by replaying the visual consequences of an eye movement back to the observer, stimulus visibility could not be used to infer the presence of an eye movement. The present experiments suggest that observers were unable to develop even a shallow understanding of how their eye movements related to seeing the stimulus. Observer’s average confidence ratings remained close to the scale’s midpoint (the point of highest uncertainty)—especially for eye movements generated in **Experiment 2** (**Fig. 3d**). Additionally, microsaccade sensitivity was overall low, suggesting that participants had limited information about their eye movement to infer how it affected stimulus visibility (**Fig. 3c**). If anything, our data suggests that participants tried to assign causality by estimating how much they lacked a sense of control over stimulus visibility. Our results indicate that observers were able to detect the absence of a causal relationship while they struggled to correctly determine when an eye movement caused the stimulus percept: In **Experiment 1**, certainty was highest when participants reported that they had *not* caused the stimulus to become visible, and we replayed a previous eye movement back to them (**Fig. 3d**). Fittingly, in trials with similarly replayed eye movements, observers were least certain when expressing the (incorrect) belief that their own microsaccade allowed for stimulus detection. In contrast, when a generated microsaccade rendered the stimulus visible, certainty ratings were not statistically different for correct and incorrect causal assignments. Observers expressed comparable levels of certainty when (correctly) claiming that their eye movement allowed for stimulus detection and when expressing the (incorrect) belief that the change stimulus visibility was not due to a microsaccade (**Fig. 3d**).

While we already mentioned low microsaccade sensitivity as one potential explanation, a second, equally interesting explanation can be found in the paradigm itself. We fundamentally broke how eye movements naturally relate to changes in retinal inputs: We presented a stimulus that is perceived during a saccade while the pre- and post-saccadic retinal images were largely identical. This contrasts with natural situations, in which the retinal input changes drastically across saccades. Replayed eye movements, on the other hand, were purely visual events—and our observers’ task, hence, much more natural. In other words, by decoupling eye movements and their visual consequences, our paradigm allowed to examine the role of intention shapes sensorimotor awareness in the active observer. However, by deviating from how eye movements and their perceptual consequences are linked in natural viewing, we obscured the relationship between a microsaccade and stimulus perception. This hindered observers’ understanding of how their eye movements influenced stimulus visibility, explaining observers’ low certainty when attributing stimulus visibility to their own eye movements. Because replaying the retinal consequences of a previous eye movement was a purely visual event (that did not break with natural viewing in the same way), observers’ certainty was much higher when correctly asserting that the stimulus became visible despite the absence of a microsaccade.

### Theoretical implications

To summarize, we found no evidence that the parameters of eye movements affected the degree to which observer were aware of them, as miniscule saccades of similar amplitude, peak velocity, and duration could either be detected (i.e., intended and unintended microsaccades) or not (i.e., spontaneous microsaccades). While our data provides some evidence that an eye movement’s visual consequences influences movement awareness (**Fig. 3c**), decoupling eye movements and their visual consequences in our paradigm revealed intention as another crucial factor for sensorimotor awareness of even the most minute of actions: We found low awareness of actions without intention (i.e., low sensitivity for spontaneous microsaccades from **Exp. 2**), and heightened awareness for actions congruent with an intention (i.e., increased sensitivity for intended microsaccades from **Exp. 1**) as well as intention-incongruent actions (i.e., increased sensitivity for unintended microsaccades from **Exp. 1**). Our data therefore support that fixating constitutes a process that is also intended and controlled much like the generation of a saccade.

### Conclusion

For this study, we developed a novel paradigm that allowed us to dissociate the role of action intention and an action’s sensory consequence for action awareness, two factors that previous research has typically confounded. Our results provide strong evidence that observers can, in principle, detect even the smallest possible eye movements. Action intention is the main driver of the perception of these tiny visual motor acts: Observers were significantly more sensitive to microsaccades when they intended to make or avoid them compared to when such microsaccades occurred spontaneously. Instantaneous sensory consequence did not lead to a similar increase in saccade sensitivity, demonstrating that sensorimotor contingencies did not enhance eye movement awareness. Taken together, our data support the conclusion that intention opens a gate to motor awareness even for unintended actions. Consequently, even microsaccades—the body’s smallest actions—can be detected, whereas these movements typically escape awareness in the absence of a related intention.

## Acknowledgements

JNK was supported by the Berlin School of Mind and Brain, Humboldt-Universität zu Berlin. SO was supported by the DFG (OH 274/4-1) and funding from the Heisenberg Programme of the DFG (OH 274/5-1). MR was supported by the European Research Council (ERC) under the EU’s Horizon 2020 research and innovation program (grant agreement no. 865715), and by the DFG (grants RO3579/8-1 and RO3579/10-1).

## STAR Methods

### RESOURCE AVAILABILITY

#### Lead contact

Information and requests regarding resources for this study should be directed to and will be fulfilled by the lead contact, Jan-Nikolas Klanke

### Materials availability

There are no restrictions for the distribution of materials.

### Data and code availability

- The preregistration, data, and all original code for **Experiment 1** has been deposited at the Open Science Framework and will be made publicly available as of the date of publication. [LINK WILL FOLLOW HERE].
- The preregistration, data, and all original code for **Experiment 2** has been deposited at the Open Science Framework and will be made publicly available as of the date of publication. [LINK WILL FOLLOW HERE].

### EXPERIMENTAL MODEL AND SUBJECT DETAILS

In **Experiment 1**, a total of 10 participants were recruited by means of the “Psychologischer Experimental-Server Adlershof” (PESA) of the Humboldt-Universität zu Berlin. Participants (6 female, 1 diverse) had a mean age of 24.2 years old (*SD* = 4.6, *min* = 18, *max* = 31), and all 10 were right-handed and 8 were right-eye dominant. All 10 participants had normal or corrected-to-normal vision. Participants were paid upon completion of the last session. The compensation was based on an hourly rate of €10/hour. Alternatively, psychology students could choose to obtain participation credit (1 credit per 15 minutes of participation) required for the successful completion of their bachelors’ program.

In **Experiment 2**, we recruited a total of 10 participants. Because we wanted a direct comparison between experiments, we tried to recruit the same participants for both experiments but were only able to successfully re-recruit 3 participants. The additional participants were recruited by means of the “Psychologischer Experimental-Server Adlershof” (PESA) of the Humboldt-Universität zu Berlin. Participants (7 female, 0 diverse) had a mean age of 26.5 years old (*SD* = 6.2, *min* = 20, *max* = 34). All participants were right-handed, right-eye dominant, and had normal or corrected-to-normal vision. Participants were paid upon completion of each experiment: Compensation was based on an hourly rate of €8/hour and a bonus payment of €4 for the completion of the final session. Psychology students could again alternatively choose to gain participation credit (1 credit per 15 minutes of participation) required for the successful completion of their bachelors’ program.

**Experiment 1** and **2** were approved by the ethics committee (Ethikkomission) of the Institut für Psychologie at the Humboldt-Universität zu Berlin and conducted in agreement with the Declaration of Helsinki (‘World Medical Association Declaration of Helsinki’, 2013) and the General Data Protection Regulation (GDPR) of the EU. All participants provided informed consent in writing before the start of the first session.

For **Experiment 1** and **2**, we pre-registered three exclusion criteria that ensured that participants would not participate if they showed the inability to execute stable fixation and correct eye movements:

- The inability to complete at least 4 blocks during the first experimental session due to fixation failures led to immediate exclusion from the experiment.
- If we could not detect more than 3.5 (**Exp. 1**) or 2.5 microsaccades (**Exp. 2**) in the crucial time window (200-800 ms re stimulus onset) of trials with generated eye movements across each block of the first session, we likewise excluded the participant from further testing.
- During data analysis, we double-checked the eye movement data offline. We excluded all participants that generated less than 150 microsaccades in the crucial time window (200-800 ms re stimulus onset) of generated microsaccade condition trials. Especially the second and third criteria were set to ensure that we would obtain enough data from each participant for the planned analyses.

In **Experiment 1**, no participant was excluded from data collection, but two participants decided against further participation after partially completing the first session and having trouble with the task and/or eye tracker. We could have excluded one participant for their overall low number of microsaccades after the completion of all trials (we were only able to detect 105 microsaccades in their data) but decided against it for economic reasons.

In **Experiment 2**, a total of six participants were excluded: four participants were excluded because they generated less than the required amount of microsaccades in the first session of the experiment, two participants because their overall number of microsaccades was vastly lower than 150. We could have excluded two more participants based on the third exclusion criterion (they generated slightly less than the required 150 microsaccades in total: 134 and 139 respectively) but decided against it for economic reasons as well. Data collection for both experiments was heavily affected by COVID-19.

## METHOD DETAILS

### Apparatus

Participants were seated in a dark room in front of a screen at a distance of 340 cm and their head stabilized using a chin rest. We projected visual stimuli on a 141.0 x 250.2 cm video-projection screen (Stewart Silver 5D Deluxe; Stewart Filmscreen, Torrance, CA, USA) using a PROPixx DLP (960 × 540 pixels; VPixx Technologies Inc., Saint Bruno, QC, Canada) with a refresh rate of 1440 Hz. We recorded participants’ eye positions of both eyes with a head-mounted eye tracker at a sampling rate of 500 Hz (EyeLink 2 Head Mount; SR Research, Ottawa, ON, Canada). The experiments were controlled on a workstation running the Debian 8 operating system, using Matlab (Mathworks, Natick, MA), the Psychophysics Toolbox 3 (Brainard, 1997; Kleiner et al., 2007; Pelli, 1997) and the EyeLink Toolbox (Cornelissen et al., 2002).

### General Methods

In **Experiment 1**, we wanted to compare intended microsaccades (i.e., eye movements generated under explicit movement instructions) and unintended (i.e., eye movements generated under explicit fixation instructions). We deployed an adapted version of a memory-guided microsaccade paradigm (Willeke et al., 2019), which presented the instructions for each trial during an initial fixation interval. During this interval, participants were either instructed to retrain their gaze position at the onscreen location indicated by the prolonged presentation of (only) the fixation point (50% of all trials), or to make an eye movement to another location specified by an eye movement target, presented in addition to the fixation point (50% of trials). The target was a white circle with the same diameter as the dot at the center of the fixation point (0.2 dva) and presented at a radial distance of either 0.2 dva, 0.4 dva, 0.6 dva, 0.8 dva, or 1 dva relative to the fixation dot. To introduce some variation to the microsaccade target location, we allowed for the microsaccade target to be displaced along the circumference of its radial distance to the fixation dot. This displacement was sampled from a normal distribution centered on 0 deg and with a standard deviation of 25 deg, resulting in onscreen locations of the microsaccade target that vary with smaller vertical than horizontal displacements.

#### Fixation-check interval

Before the start of each trial, a target-shaped central fixation point appeared before an otherwise grey background. The fixation dot (inner part) had a diameter of 0.2 dva while the outer ring had a diameter of 0.6 dva. Before the onset of each trial, a fixation control routine was run that required the gaze position of the observer to be inside a circular region (3 dva in diameter) around the fixation point. The trial began when the fixation control was successful for at least 200 ms. The fixation point appeared at the onscreen location on which the stimulus presentation was centered in the following.

#### Fixation interval

The fixation interval (present only in **Experiment 1**) started as soon as the outer ring of the fixation point disappeared. The interval duration varied randomly between 400 ms and 500 ms to avoid routine anticipatory eye movements. Fixation dot and microsaccade target remained visible for the entire duration of the fixation interval. Participants were instructed to keep their gaze locked on the fixation dot without making any eye movements as long as the dot was visible (i.e., for the duration of the fixation interval) in instructed fixation as well as instructed eye movement condition trials. If a microsaccade target was displayed additionally, participants were to memorize the onscreen location of the target and generate an eye movement to this location as soon as the fixation dot and microsaccade target disappeared (i.e., in the beginning of the stimulus presentation interval). In trials without microsaccade target presentation, participants were instructed to keep their eye position centered on the location of the fixation dot even after it disappeared.

#### Stimulus presentation interval

The disappearance of the fixation point (inner part and outer part) indicated the start of the stimulus presentation interval. Stimulus presentation lasted for 1000 ms independent of condition. The position of the stimulus was determined randomly in each trial, but its midpoint was always within ±4 dva relative to the screen center (horizontally as well as vertically). Between the stimulus presentation and the response interval, there was a short delay of 50 ms during which nothing was presented on the gray screen.

#### Response interval

In the response interval, participants had to answer two simple yes-no questions and, depending on their response to these, a confidence rating. At first, we displayed the question “Did you perceive a stimulus flash?” on the screen. Participants could respond with either ‘Yes!’ or “No!’ (both response options were presented onscreen below the question as well). In a second step, participants had to indicate whether they believe to have generated an eye movement. To this end, we displayed the question “Do you think you generated an eye movement?” together with the two response options from before. In both cases, responses were submitted by pressing either the left or the right arrow (i.e., the arrow key in the direction of the chosen response option).

Participants’ responses to these first two questions determined the presentation of the final stage of the response phase: If participants reported that they perceived a stimulus flash and that they thought they generated an eye movement, they were be asked: “How sure are you that the stimulus was caused by an eye movement?”. If they report to have perceived the stimulus flash but that they did not generate an eye movement, the question instead was: “How sure are you that the stimulus flash was not caused by an eye movement. To respond to this final question, participants had to choose one of four options displayed on a continuous scale: “not sure”, “rather not sure”, “rather sure”, and “very sure”. Participants selected their response by adjusting the position of a response prompt via the left and right arrow keys. If the response prompt assumes the desired position, participants logged-in their answers by pressing the space key. The lateralization of the response options (i.e., which option is displayed on which side of the stimulus center) remained the same for all sessions of one participant but was counterbalanced between participants.

### Variations in Experiment 2

In **Experiment 2**, the fixation check interval, stimulus presentation interval, and response intervals were the same as in **Experiment 1**. Because we wanted to compare intended and unintended microsaccades from **Experiment 1** to spontaneous microsaccades, we removed the fixation interval before the stimulus presentation. Participants were informed that some microsaccades occurred spontaneously before the start of the first session and were informed that trials would abort if their gaze position deviated too much from the location indicated during the fixation check interval but received no further instruction regarding their eye movement behavior.

### Online control of eye positions

During **Experiments 1** and **2**, participants’ eye positions were tracked. Eye and screen coordinates were aligned by conducting standard nine-point calibration and validation procedures before the first trial of each session and whenever necessary. Blinks and deviations in gaze position (>1.5 dva from fixation) were likewise monitored in both experiments and led to an abortion of the trial. Aborted trials were repeated at the end of each block in randomized order.

### Pre-processing

Binocular microsaccades were detected using an algorithm described by Engbert and Mergenthaler (2006) in **Experiment 1** and **2**. For the velocity threshold, we used a λ of 5 and minimum microsaccade duration of 6 ms (3 data samples). To exclude potential over- or undershoot corrections, two microsaccades were merged if the interval between them was shorter than 10 ms (5 data samples).

For the replay of the retinal consequence of a microsaccade, we used the gaze positions of the dominant eye of each observer recorded during binocularly detected microsaccades. To allow for a direct comparison between the visibility of the stimulus between conditions, we only replayed the retinal consequences of microsaccades that were recorded in the crucial time window (200-800 ms after stimulus onset) of trials in which a generated microsaccade could render the stimulus visible. Additionally, we did not use the raw microsaccade data for the stimulus but pre-processed the recorded gaze trajectories. In a first step, we re-centered the recorded gaze positions of each microsaccade on the origin by subtracting the coordinates of the first data sample form all remaining samples. In a second step, we excluded all gaze positions sampled after the microsaccade reaches its maximum amplitude because microsaccades frequently follow a curved trajectory that would likely lead to blurry or obscure percepts when replayed. If a microsaccade was shorter than 6 ms (3 data samples) before its maximum amplitude was reached, it was excluded altogether. In a third step, we projected the recorded eye positions onto the saccade vector by recalculating the location of each gaze position during the saccade relative to its amplitude. In step number four, we fit a gamma function to the velocity profile of the saccade vector. Optimal fits were determined by means of a root mean square error (RMSE) procedure. We additionally ensured the quality of the fits by excluding microsaccades for which the root mean square error deviated more than two standard deviations from the mean of all RMSEs of the same session from one participant. In step number five, we redistributed the gaze positions along the saccade vector based on the fitted velocities. To compensate for the difference between the frequency of the eye tracker (500 Hz) and the refresh rate of the projector used for the display (1440 Hz), the recalculation of the gaze position along the saccade vector was combined with an upsampling mechanism that padded the number of data points along the saccade vector according to the fitted velocity profile. In a sixth step, we checked that the upsampling mechanism did not lead to velocity profiles that were biologically implausible. To this end, we excluded microsaccades for which the peak velocity of the upsampled saccade vector was three times higher (or more) than the peak velocity as predicted for microsaccades of maximum amplitude (i.e., 1 dva) by the main sequence (a known curvilinear relationship between saccade amplitude and peak velocity, see Zuber et al., 1965) of the individual participant. In a final step, we inverted the coordinates of the upsampled saccade vector: We wanted to replay the retinal consequences of the image during a microsaccade back to the observers, and the retinal image is always shifts in the opposite direction of the eye movement. The same data preprocessing was used in **Experiment 1** and **2**.

### Exclusion of trials from analyses

Because the replay differed between the first and later sessions of each participant, data obtained in session one were not considered in the main analysis. Trials in which a replayed microsaccade could render the stimulus visible and in which the participant generated at least one (additional) microsaccade as well as trials in which the participant generated more than one microsaccade in the stimulus presentation interval were likewise excluded. Note that trials with accidental microsaccade generation in the fixation interval of **Experiment 1** were not excluded: Participants were instructed to try and make an accurate eye movement again at the beginning of the stimulus presentation interval (i.e., after the disappearance of fixation point and saccade target) and only report for those eye movements.

We also disregarded trials with generated microsaccades larger than 1 dva and when the microsaccade failed to occur in the crucial time window of the stimulus (200-800 ms re stimulus onset). Finally, due to an error in the code, **Experiment 2** included the replay of microsaccades detected only monocularly. To account for this mistake, we excluded trials in which erroneously detected microsaccades were replayed from all analysis. For an overview over the number of valid trials per experiment, eye movement and stimulus condition, see ***Table 1***).

**Table 1:**
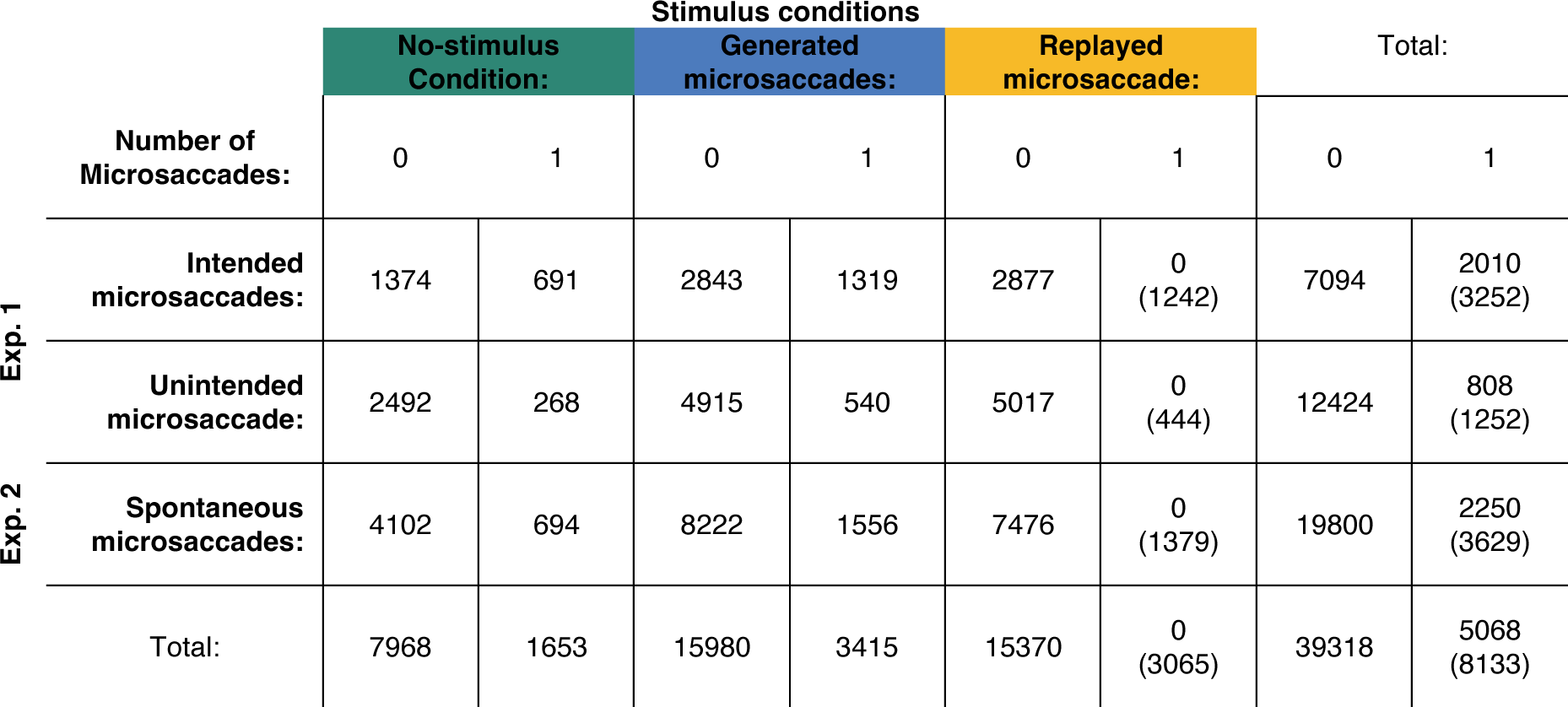
Overview over the number of valid trials in each stimulus display and eye movement condition.

## QUANTIFICATION AND STATISTICAL ANALYSIS

### Motor control for microsaccades

#### Saccade rates

To investigate motor control for microsaccades, we calculated individual saccade rates separately for different eye movement types: intended and unintended microsaccades (**Exp. 1**), as well as spontaneous microsaccades (**Exp. 2**). Saccade rates were calculated individually as the number of trials with a saccade divided by the number of all trials per participant (irrespective of stimulus condition). To assess how well participants could adept their instructed eye movements to the different target distances (ranging from 0.2 to 1 dva), we further categorized trials with intended microsaccades according to those distances. We predicted higher rates when participants were instructed to move their eyes (i.e., for intended microsaccades, **Exp. 1**), compared to when they were instructed to fixate (i.e., unintended microsaccades, **Exp 1**), or when they received no instruction (spontaneous microsaccades, **^Exp. 2^**^; rate^_spontaneous_ ^≅ rate^_unintended_ ^< rate^_intended_^).^

To determine if observers generated more saccades when instructed to do so in **Experiment 1**, we calculated a paired two-sided t-test for the within-subject comparison of intended and all unintended microsaccades. To assess the effect of target distance, we calculated a one-way repeated-measures analysis of variance (rmANOVA) on saccade rates with intended microsaccades, categorized based on target distance. To compare saccade rates between experiments, we employed two two-sided independent samples t-tests, compararing average rates of spontaneous microsaccades (**Exp. 2**) and unintended microsaccades (**Exp. 1**), as well as spontaneous (**Exp. 2**) and intended eye movements (**Exp. 1**).

#### Saccade amplitudes

Because we were interested in motor control for intended microsaccades, we calculated average amplitudes per participant and the five different target distances (0.2–1 dva). We did not pre-register specific hypotheses but would predict larger saccade amplitudes in trials with greater target distance.

To determine if greater target distances indeed led to saccades with larger amplitudes, we fit a linear mixed effects model to the unaggregated intended microsaccades from **Experiment 1.** The model predicted saccade amplitude with target distance by using a restricted maximum likelihood (REML) method. Participants were included in the model as random effects.

### Visual sensitivity to intra-saccadic stimulation

#### Eye movement generation

We examined observer’s visual stimulus sensitivity based on their responses to the first question “Did you perceive a stimulus flash?”. We calculated individual hit rates based on the number of positive “Yes!”-responses in trials with stimulus. Similarly, individual false alarm rates were calculated based on “Yes!”-responses in trials without stimulus (no-stimulus condition trials). Because false alarm reports were very rare (40% of our participants did not report a single false alarm, and those who did only reported 1.57 false alarms on average), we decided not to calculate separate rates depending on saccade generation (as pre-registered), but combined rates for trials with and without eye movements. To determine if stimulus visibility depended on the type of eye movement generated, we determined different rates for trials with small intended and unintended saccades (**Exp. 1**), as well as spontaneous microsaccades (**Exp. 2**). To determine visual sensitivity per participant and condition, individual hit and false alarm rates were z-transformed and subtracted (i.e., d^+^ = z(Hits) − z(FAs)). We predicted that, while visual sensitivity should depend on eye movement generation (i.e., d’_0 MS_ < d’_1 MS_), it should not differ between generated and replayed microsaccades (i.e., d’_generated_ ≅ d’_replayed_). Similarly, we did not expect visual sensitivity to differ based on eye movement type (i.e., ^d’^_intended_ ^≅d’^_unintended_ ^≅ d’^_spontaneous_).

To determine if the stimulus was invisible during stable fixation, we calculated averaged sensitivity indices per eye movement type and compared their corresponding 95% confidence intervals (*CI_95%_*) against 0. Significant differences between eye movement types were determined by calculating a paired two-sided t-test for the within-subject comparison of intended and unintended eye movements (**Exp. 1**), and two-sided independent samples t-test for the comparison between combined intended and unintended microsaccades (**Exp. 1**) and spontaneous microsaccades (**Exp. 2)**. The effect of eye movement generation in **Experiment 1** was determined by a two-way rmANOVA with visual sensitivity indices as the depended variable, and the stimulus condition (generated vs. replayed) and eye movement type (intended vs. unintended) as within-subject factors. To compare eye movements from different experiments, we calculated a mixed-measures ANOVA with stimulus condition (generated vs. replayed) as a within-subject factor and experiment (**Exp. 1** vs **Exp. 2**) as between-subject factor.

#### Eye movement kinematics

Because visual sensitivity should depend on the degree of retinal stabilization, we additionally calculated retinal velocity of the stimulus. To obtain retinal velocities, we subtracted the constant speed of the phase shift from the peak velocity of each microsaccade. We used directed speeds, i.e., positive values for rightward and negative values for leftward oriented phase shifts or saccade directions. Retinal velocity of 30 dva/s or less were labelled ‘low’, velocities surpassing 30 dva/s were labelled ‘high’. We calculated hit and false alarm rates as well as visual sensitivity separately for generated and replayed eye movements of all three types (**Exp. 1**: intended and unintended microsaccades; **Exp. 2**: spontaneous microsaccades) and according to the resulting retinal velocity of the stimulus (for more details see previous paragraph). We predicted that a higher retinal stability of the stimulus would lead to increased visual sensitivity. Consequently, microsaccades (irrespective of type) that lead to lower retinal velocities of the stimulus should yield higher sensitivity compared to trials with higher retinal velocity (i.e.,d’_high vel._ < d’_low vel._).

To determine if lower retinal stimulus velocities indeed led to higher visual sensitivity for eye movements from **Experiment 1**, we calculated a three-way rmANOVA with the within-subject factors retinal velocity (low vs. high velocity), stimulus condition (generated vs. replayed), and eye-movement type (intended vs. unintended). To compare these results to eye movements in **Experiment 2**, we calculated a three-way two-way mixed-measures ANOVA with the within-subject factors retinal velocity (low vs. high velocity) and stimulus condition (generated vs. replayed), and the factor experiment (**Exp. 1** vs. **Exp. 2**) as a between-subject factor. Significant differences between factors were determined by calculating paired two-sided t-test for within-subject comparisons or two-sided independent samples t-test to compare between experiments whenever necessary.

One participant was excluded from this analysis, because a hit rate in a particular condition could not be computed (i.e., unintended microsaccades that led to a high retinal velocity of the stimulus).

### Eye movement sensitivity

#### Stimulus presentation

To determine how sensitive participants were towards their own eye movements, we analyzed participants responses to the second question of the response phase: “Do you think you generated an eye movement?”. To gain a better understanding of how eye movement awareness was affected by stimulus presentation, we calculated sensitivity separately for stimulus present and absent trials. Hit rates in stimulus present trials were calculated based trials with generated eye movement for which participants correctly reported believing to have generated a microsaccade. False alarm rates were, conversely, calculated based on trials with replayed microsaccades (i.e., in the absence of a generated saccade) for which participants incorrectly reported the same belief. In stimulus absent trials, hits and false alarm rates were calculated identically with the only difference that trials were taken solely from the no-stimulus condition. To assess if awareness differed between different types of eye movements, we additionally categorized trials by eye movement type; intended (**Exp. 1**), unintended (**Exp. 1**), and spontaneous microsaccades (**Exp. 2**). We expected low sensitivity towards spontaneous and unintended microsaccades, but increased sensitivity towards intended eye movements (i.e., d’_spontaneous_ ≅ d’_unintended_ < d’_intended_). No predictions about stimulus presentation were preregistered, however, we expected that—because the stimulus was presented saccade-contingently—stimulus presentation would facilitate detection for all types of microsaccades (i.e., d’_absent_ < d’_present_).

To determine eye movement sensitivity in **Experiment 1**, we calculated a two-way rmANOVA with the within-subject factors eye movement type (intended vs. unintended) and stimulus presence (present vs. absent). Spontaneous microsaccades from **Experiment 2** were compared to results from the first experiment by calculating a two-way mixed-measures ANOVA with the within-subject factor stimulus presentation (absent vs. present) and the between subject factor experiment (Exp. 1 vs. Exp. 2). Paired two-sided t-test for within-subject comparisons or two-sided independent samples t-test to compare between experiments were calculated to determine significance whenever necessary.

#### Causal assignment

To investigate if observers were able to detect whether their eye movements caused the high-temporal frequency stimulus to become visible, we analyzed their responses in the final part of the response phase. In this phase, we displayed one of two questions, depending on their previous responses: “Do you think your eye movements caused the stimulus flash?” if a participant had reported the presence of an eye movement, and “Do you think your eye movements did not cause the stimulus flash?” when they reported no eye movement. Unlike before, participants could respond on a 4-point scale, spanning form “very sure” to “very unsure”. While, according to our pre-registration, we planned to use this response schema to calculate meta-d’, we ultimately decided that our data could be better understood by a simpler analysis: We assigned each response option a fixed score between -1.5 and 1.5 depending on the level of certainty (i.e., 1.5 for “very sure”, 0.5 for “rather sure”, -0.5 for “rather unsure”, and -1.5 for “very unsure”), before calculating average scores per stimulus condition and participants. Because we expected that participants ability to correctly assign causality to depend on eye movement awareness and because participants were similarly sensitivity towards their intended and unintended microsaccades in **Experiment 1**, we decided to neglect differences between these two types of saccades and categorized eye movements only by experiment for this analysis to increase its power.

We predicted that the ability to assign causality correctly directly depended on participants’ eye movement awareness. Because eye movement sensitivity was higher in the first compared to the second experiment, we expected higher scores for **Experiment 1** compared to **Experiment 2** (i.e., scores_exp.2_ < scores_exp.1_).

To analyze if observers assigned causality correctly in **Experiment 1**, we calculated a two-way rmANVOA using certainty scores as the dependent variable and stimulus condition (generated vs. replayed) and correctness of the assignment (correct vs. incorrect) as within-subject factors. For **Experiment 2**, we replicated the analysis with an identical two-way rmANOVA deploying the within-subject factors stimulus condition (generated vs. replayed) and correctness of the assignment (correct vs. incorrect) again. Paired two-sided t-test for within-subject comparisons or two-sided independent samples t-test to compare between experiments were calculated to determine significance whenever necessary. One participant was excluded in **Experiment 1**, another participant was excluded from **Experiment 2**—both were excluded because we could not calculate certainty for trials with a microsaccade was generated but none reported.

### Supplements

#### Microsaccade rates

To better understand generation of intended microsaccade over small target distances (**Exp. 1**), we compared rates for successive distances via paired t-test (all *p*-values reported here are Bonferroni-corrected to adjust for multiple comparisons).

While we found significant differences in rates for smaller target distances (0.2 dva vs. 0.4 dva: *t* (9) = –3.27, *p* = 0.039; 0.4 dva vs. 0.6 dva: *t* (9) = –3.15, *p* = 0.039). We did not observe significant differences for comparisons over larger distances (0.6 dva vs. 0.8 dva: *t* (9) = –0.17, *p* > 0.250; 0.8 dva vs. 1.0 dva: *t* (9) = 0.57, *p* > 0.250). This supports the earlier conclusion that saccade rates increase with increasing target distances.

In a second step, we compared saccade rates of intended microsaccades for each target distance to unintended microsaccades by again conducting two-sided paired t-tests for each comparison (with Bonferroni-correction for multiple comparisons). Our testes revealed insignificant differences between the rates of unintended and intended microsaccades when the target distances were smaller or equal to 0.4 dva (unintended vs. 0.2 dva: *t* (9) = 1.64, *p* > 0.250; unintended vs. 0.4 dva: *t* (9) = 2.91, *p* = 0.086). When target distances exceeded 0.4 dva, however, the intended and unintended microsaccade rates differed increasingly significantly (unintended vs. 0.6 dva: *t* (9) = 3.30, *p* =0.046; unintended vs. 0.8 dva: *t* (9) = 3.78, *p* = 0.022; unintended vs. 0.8 dva: *t* (9) = 4.17, *p* = 0.012).

Lastly, comparing between spontaneous (**Exp. 2**) and intended microsaccades generated over the different target distances reveals no significant difference—irrespective of target distance (all Bonferroni-correct *p*s > 0.250).

Taken together, our finding suggests that task difficulty to reliably generate intentional microsaccades increases when target distances get smaller—particularly when eye movements must be generated without foveal anchor. Interestingly, insignificant differences between microsaccade rates over small target distances and unintentional microsaccades suggest that participants perform at a level compared to intended fixation when trying to make eye movements smaller than 0.4 dva by memory. Of course, this result has to be interpreted with caution, as the smallest saccades are also the hardest to detect with video-based eye tracking equipment.

#### Accuracy and precision of intended microsaccades

To investigate motor control for intended microsaccades from **Experiment 1**, we calculated averaged accuracy and precision of eye movements over the five target distances (ranging from 0.2 to 1 dva) for each participant. We determined significance by calculating one-way rmANOVAs with precision or accuracy as the depended variable and target distances (ranging from 0.2 to 1 dva) as within-subject factor.

We found that while our participants tended to overshoot when target distances were small (0.2 dva: 0.22±0.09, 0.4 dva: 0.10±0.11), accuracy was high for the medium distance (0.6 dva: –0.04±0.11). Conversely, for longer target distances, participants tended to undershoot the target (0.8 dva: –0.16±0.10; 1.0 dva: –0.35±0.12; **Fig. S1a**). Unsurprisingly, a one-way rmANOVA revealed a significant effect of target distance on saccade accuracy (*F* (4,36) = 106.22, *p* < 0.001). While observers did adapt saccade amplitudes to the target amplitudes (see section *Motor control for microsaccades* in results), the pattern of overshooting eye movements over smaller target distances and undershooting eye movements over larger target distances reveals a preference to generate microsaccades of medium size.

Precision on the other hand, is near identical over all target distances (0.2 dva: 0.18±0.05; 0.4 dva: 0.18±0.04; 0.6 dva: 0.18±0.03; 0.8 dva: 0.18±0.04; 1.0 dva: 0.21±0.04; **Fig. S1b**), indicating that saccades were executed with equal precision irrespective of target distance. We conducted a one-way rmANOVA precision as depended variable and target distance as within-subject factor to corroborate this finding (*F* (4,36) = 1.23, *p* > 0.250).

**Figure S1.**
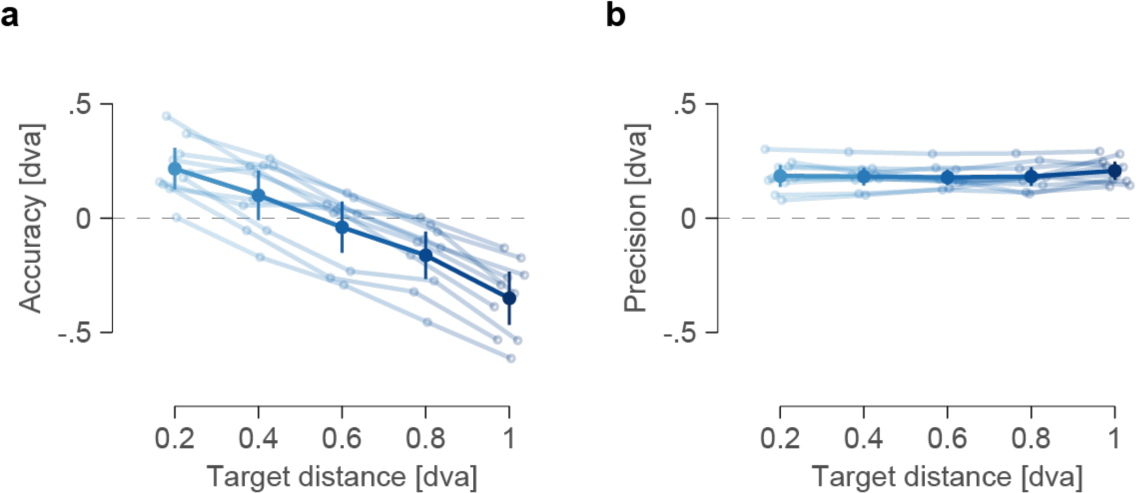
Intended microsaccades overshot small distances, undershoot long distances, but are executed with precision regardless. **a** Accuracy and **b** precision of intended microsaccades from Experiment 1. In all panels, small circles indicate individual observers’ means, filled dots represent sample means. Lines connect dots of individual participants. Error bars indicate 95% confidence intervals.

#### Parameters of different eye movement types

To determine similarities and differences between the different types of eye movements, we compared four different sets of parameters for intended, unintended (**Exp. 1**), and spontaneous microsaccades (**Exp. 2**): amplitude, peak velocity, duration, and latency.

In this analysis, we computed the means of individual eye movement parameters for each participant before conducting comparisons between the different eye movement types. We utilized two-alternative, paired t-tests when comparing eye movement from **Experiment 1**, while for the comparison between **Experiment 1** and **2**, we employed two-alternative, between-subject t-tests.

Starting with the parameter amplitude, we found the largest amplitudes for intended microsaccades (0.56±0.10), while unintended microsaccades were markedly smaller (0.30±0.06). The difference between the two eye movement types was highly significant (*t* (9) = 7.40, *p* < 0.001). Spontaneous microsaccades had an intermediary size (0.40±0.09; **Fig. S2a**). Comparing between experiments, we found spontaneous and unintended microsaccades to be more similar in size (*t* (16.4) = –2.11, *p* = 0.051) compared to intended microsaccades (*t* (17.9) = 2.77, *p* = 0.01).

Turning to peak velocity next, we observed that intended microsaccades yielded the highest peak velocities (57.27±8.83). Unintended microsaccades, on the other hand, were characterized by significantly lower peak velocities (35.07±6.15; *t* (9) = 7.22, *p* < 0.001). Peak velocities of spontaneous microsaccades were, again, on an intermediate level (40.38±8.26; **Fig. S2b**)—matching the peak velocity of unintended microsaccades more closely (*t* (16.6) = –1.17, *p* > 0.250) than that of intended ones (*t* (17.9) = 3.16, *p* = 0.005).

Next, we investigated how the durations of eye movements differed between intended unintended, and spontaneous microsaccades. As before, our data revealed intended eye movements to have the longest durations (22.15±2.46)—particularly compared to unintended (16.76±2.07) but also spontaneous microsaccades (19.37±2.41; **Fig. S2c**). Here, only the differences between intended and unintended microsaccades turned out to be significant (*t* (9) = 8.49, *p* < 0.001), while both comparisons between experiments remained insignificant (unintended vs spontaneous: *t* (17.6) = –1.86, *p* = 0.080; intended vs. spontaneous: *t* (18.0) = 1.82, *p* = 0.084), indicating that the duration of unintended microsaccades was even shorter than that of spontaneous ones.

Lastly, we looked at saccade latencies: We found the shortest latencies for intended microsaccades (375.99±54.84), with slightly longer latencies for unintended (465.12±44.92) and spontaneous microsaccades (476.51±36.03; **Fig. S2d**). Significant differences emerged when comparing intended and unintended microsaccades (*t* (9) = –3.18, *p* = 0.011) as well as intended and spontaneous microsaccades (*t* (15.6) = –3.47, *p* = 0.003)—while the comparison of unintended and spontaneous microsaccades remained insignificant (t (17.2) = –0.45, *p* > 0.250).

**Figure S2.**
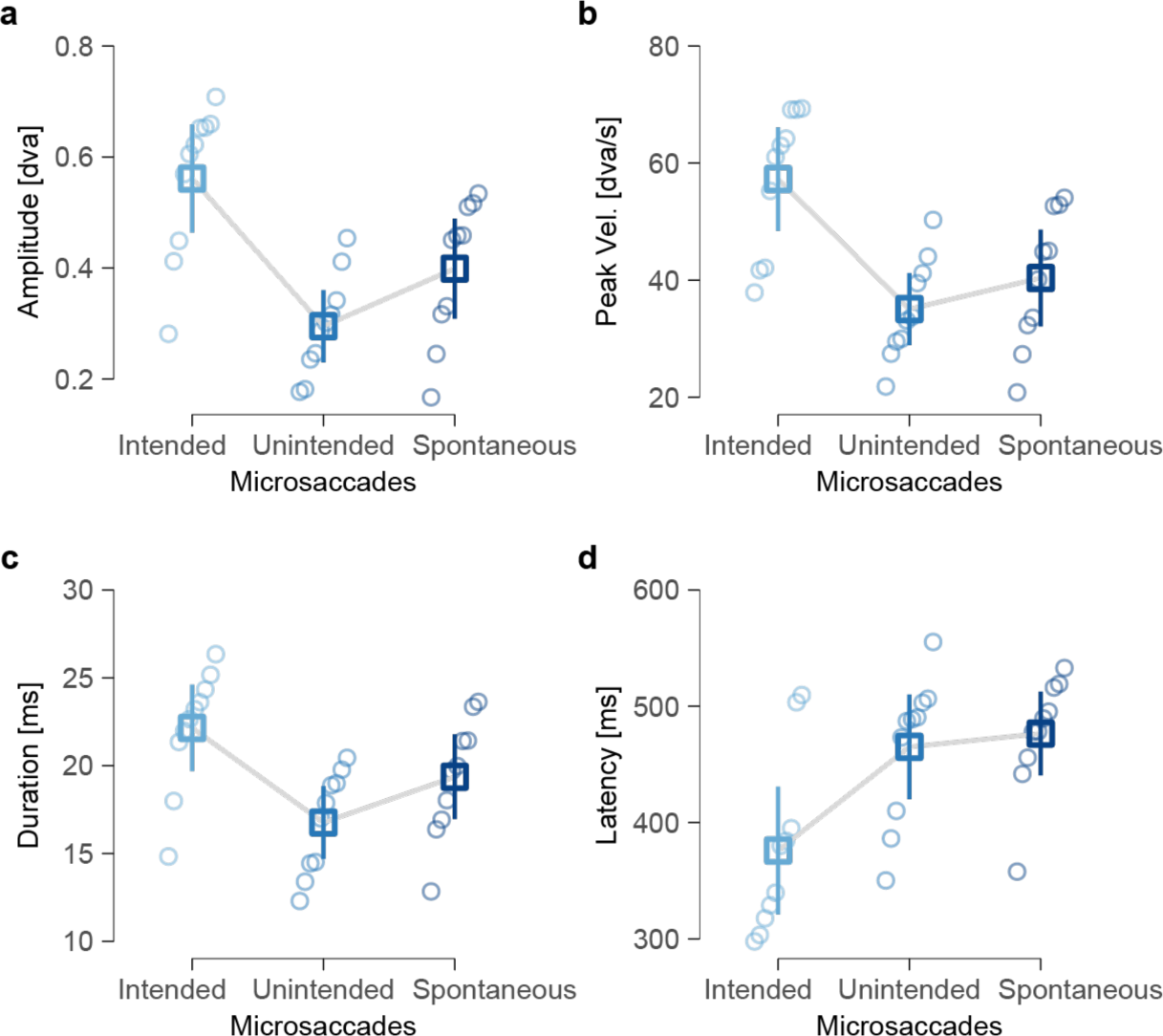
Unintended microsaccades are more similar to spontaneous than intended ones. **a** Comparison of amplitudes between intended, unintended (experiment 1), and spontaneous microsaccades (experiment 2). **b** Comparison of peak velocities between eye movement types (same as in a). **c** Comparison of duration between eye movement types (same as in a). **d** Comparison of latencies between eye movement types (same as in a). In all plots; small circle indicate individual observers’ means, squares represent sample means. Error bars indicate 95% confidence intervals.

#### Microsaccade sensitivity as a function of stimulus perception (instead of presentation)

To corroborate the dependence of sensitivity on eye movement type and determine the importance of the effect of stimulus presentation, we repeated our analysis of microsaccade sensitivity with perceptual reports of the stimulus instead of stimulus presentation (i.e., stimulus perceived vs. not perceived instead of stimulus present vs. absent).

To determine significance in **Experiment 1**, we calculated a two-way rmANOVA with eye movements sensitivity as the dependent variable. Eye movement type (intended vs. unintended) and perceptual reports (seen vs. not-seen) were included as within-subject factors. Comparison between experiments were done with a two-way mixed-measures ANOVA that, again, used eye movement sensitivity as the dependent variable and perceptual report (seen vs. not-seen) as within-subject factor and experiment (**Exp. 1** vs. **Exp. 2**) as between-subject factor. Paired two-sided t-test or two-sided independent samples t-test were calculated to determine significance whenever necessary as before.

For **Experiment 1**, we found that observers were sensitive towards intended (d’ = 0.60±0.47), and unintended microsaccades (d’ = 0.84±0.40; **Fig. S3a**) and a two-way rmANOVA revealed no significant difference between the types of eye movements (intended vs. unintended; *F* (1,9) = 0.69, *p* > 0.250). The perception of the stimulus had no significant influence in this analysis (stimulus perceived vs. not perceived; *F* (1,9) = 2.31, *p* = 0.163), indicating that observers’ microsaccade sensitivity remained unaffected irrespective of whether they perceived the stimulus. The interaction of eye movement and stimulus percept remained insignificant as well (*F* (1,9) = 0.45, *p* > 0.250).

Comparing these results to **Experiment 2**, we found that participants were less sensitive to spontaneous microsaccades, irrespective of whether they reported to have seen the stimulus (d’ = 0.22±0.28) or not (d’ = 0.16±0.14; **Fig. S3a**). Predictably, a two-way mixed-measures ANOVA revealed that microsaccade sensitivity only differed significantly when comparing eye movements from different experiments (*F* (1,18) = 11.61, *p* = 0.003), while stimulus perception failed to have a significant effect on sensitivity (stimulus perceived vs. not perceived; *F* (1,18) = 2.39, *p* = 0.139). The interaction of experiment and stimulus perception remained insignificant as well (*F* (1,18) = 0.59, *p* > 0.250).

Taken together our results suggest that the parameters of unintended microsaccades more closely resemble those of spontaneous microsaccades than intended eye movements.

**Figure S3.**
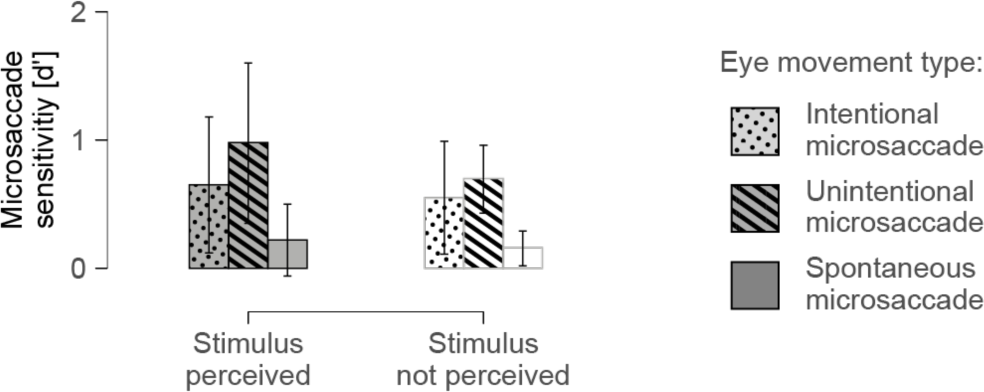
Microsaccade sensitivity based on perceptual reports.

#### Microsaccade (mis-) detection based on stimulus condition and perceptual report

Here we report the proportion of correctly detected eye movement (hits) as well as mis-detected eye movements (false alarms) for intended, unintended (**Exp. 1**), and spontaneous microsaccades (**Exp. 2**). Eye movement reports are further split according to the visual stimulus condition (generated saccade, replayed saccade, and stimulus absent) and perceptual report of the visual stimulus (stimulus perceived and stimulus not perceived). False alarm rates could not be calculated for trials with generated microsaccades for which observers reported having perceived the stimulus as reports of the stimulus as it was impossible to see the stimulus in the absence of a microsaccade and false alarms of the stimulus were exceedingly rare. We equally failed to calculate hit and false alarms in stimulus absent trials in which a stimulus percept was reported for the same reason.

We examined detection of intended and unintended microsaccades (**Exp. 1**) for generated and replayed microsaccades in relation to their perceptual consequences first. To this end, we calculated a three-way rmANOVA with hit rates as the depended variable and the within-subject factors eye-movement type (intended vs. unintended), stimulus condition (generated vs. replayed), and perceptual reports (perceived vs. not perceived) as the independent variables. The rmANOVA revealed a significant main effect of eye movement (intended vs. unintended: *F* (1,8) = 136.70, *p* < 0.001) indicating that intended eye movements (hits = 0.86±0.06) were detected significantly more often than unintended ones (hits = 0.24±0.15; **Fig. S4**). We additionally found a significant effect of perceptual report (stimulus perceived vs. not perceived: *F* (1,8) = 8.75, *p* = 0.018), indicating that eye movements were additionally detected more often in trials in which a stimulus was perceived (hit = 0.79±0.07) compared to when it was not (hits = 0.72±0.08; **Fig. S4**). The main effect of stimulus condition was not significant (generated vs. replayed: *F* (1,8) = 0.56, *p* > 0.250) and neither was any interaction (all *p*s > 0.08). One participant was excluded from this analysis because a lack of trials with unintended generated eye movement for which no stimulus percept was reported.

Comparing saccade detection rates between experiments, we calculated a three-way rmANOVA with the within-subject factors stimulus conditions and perceptual report, as well as the between-subject factor experiment (**Exp. 1** vs. **Exp. 2**) next. We found a significant main effect of perceptual report (perceived vs. not perceived: *F* (1,18) = 11.34, *p* = 0.003) and a significant main effect of experiment (**Exp.1** vs. **Exp. 2**: *F* (1,18) = 34.50, *p* < 0.001), while the main effect of stimulus presentation and all interactions remained insignificant (all *p*s > 0.059). Post-hoc comparisons revealed that—despite the low detection rates for unintended microsaccades from our first experiment—combined detection rates were still significantly higher in experiment one compared to the second experiment (*t* (12.7) = 5.90, *p* < 0.001; **Exp. 1:** hits = 0.75±0.06; **Exp. 2**: hit = 0.36±0.13; **Fig. S4**). Additional post-hoc comparisons for the significant main effect perceptual report replicate the findings from our previous analysis that trials in which a stimulus was perceived (hit = 0.68±0.11) led to significantly higher detection rates (not perceived: hits = 0.50±0.15; **Fig. S4**), even when spontaneous microsaccades were considered.

We analyzed false alarms in a separate analysis. Starting with trials from **Experiment 1** in which participants reported not having perceived a stimulus, we calculated a two-way rmANOVA with false alarm rates as the dependent variable and the within-subject factors stimulus condition (generated vs. replayed) and eye movement types (intended vs. unintended) as the dependent variables. The test revealed a significant effect of eye movement (intended vs unintended: *F* (1,9) = 86.10, p < 0.001), while stimulus condition (generated vs replayed: *F* (1,9) = 0.37, *p* > 0.250) and their interaction (*F* (1,9) = 1.20, *p* > 0.250) remained insignificant. Post-hoc comparisons revealed that false alarms are reported significantly more often for intended compared to unintended microsaccades (*t* (9) = 9.27, *p* < 0.001; intended: FAs = 0.75±0.18; unintended: FAs = 0.04±0.02; **Fig. S4**), indicating that intending to generate an eye movement increases the likelihood of reporting successful eye movement generation even in the absence of a microsaccade. Repeating this analysis for the comparison between experiments reproduced the same results: A two-way mixed measures ANOVA indicated that false alarm rates only differ when comparing eye movements between experiments (Exp. 1 vs Exp. 2: *F* (1,18) = 22.61, *p* < 0.001) not when comparing stimuli conditions with the experiments (generated vs replayed: *F* (1,18) = 0.14, *p* > 0.250; interaction: *F* (1,18) = 2.03, *p* = 0.172; **Fig. S4**): Observers were more likely to falsely report successful eye movement generation when intending to make (or suppress) an eye movement (**Exp. 1**: FAs = 0.63±0.15) compared to when they did not (**Exp. 2**: FAs = 0.18±0.14), supporting our previous supposition that false alarm rates were increased because of observers’ intention to saccade—facilitated only in **Experiment 1**.

Lastly, to investigate how seeing the stimulus affected false alarms, we calculated a two-way rmANOVA with the factors eye movement type (intended vs. unintended) and stimulus report (perceived vs. not perceived) for replayed eye movements only (since false alarms depending on stimulus perception are distributed equally only for replayed eye movements). We again found a significant main effect of eye movement (*F* (1,9) = 97.90, *p* < 0.001), indicating a much higher false alarm rate for intended (FAs = 0.77±0.16) than for unintended microsaccades (FAs = 0.07±0.04; **Fig. S4**). In addition, the factor perceptual report was significant as well (*F* (1,9) = 6.60, *p* = 0.030), with slightly higher false alarm rates for trials in which a stimulus was perceived (FAs = 0.67±0.12) than trials in which it remained imperceptible (FAs = 0.62±0.16). However, a two-alternative post-hoc t-test revealed this comparison to be marginal (*t* (9) = 2.21, *p* = 0.054; **Fig. S4**). The interaction between eye movement type and perceptual report remained insignificant (*F* (1,9) = 0.003, *p* > 0.250).

We again found the same results when comparing between experiments: A two-way mixed-measures ANOVA reveled a significant effect of the between-subject factor experiment (**Exp. 1** vs **Exp. 2**: *F* (1,18) = 15.91, *p* = 0.001; **Fig. S4**), indicating that observers misreported an eye movement more often in our first experiment (FAs = 0.65±0.14) compared to **Experiment 2** (FAs = 0.30±0.14). We also found a significant effect of perceptual report (perceived vs not perceived: *F* (1,18) = 9.53, *p* = 0.006; **Fig. S4**), with significantly higher false alarm rates in trials in which observers saw the stimulus (*t* (19) = 2.89, *p* = 0.009; perceived: FAs = 0.55±0.12; not perceived: FAs = 0.40±0.14). We found no interaction between experiment and stimulu perception (*F* (1,18) = 3.70, *p* = 0.070).

Taken together, our analyses suggest that it is neither hits, nor false alarms alone that result in a similar sensitivity for intended and unintended microsaccades. Instead, it is their shared ratio of hits to false alarms that produces the effect reported in the results section.

**Figure S4.**
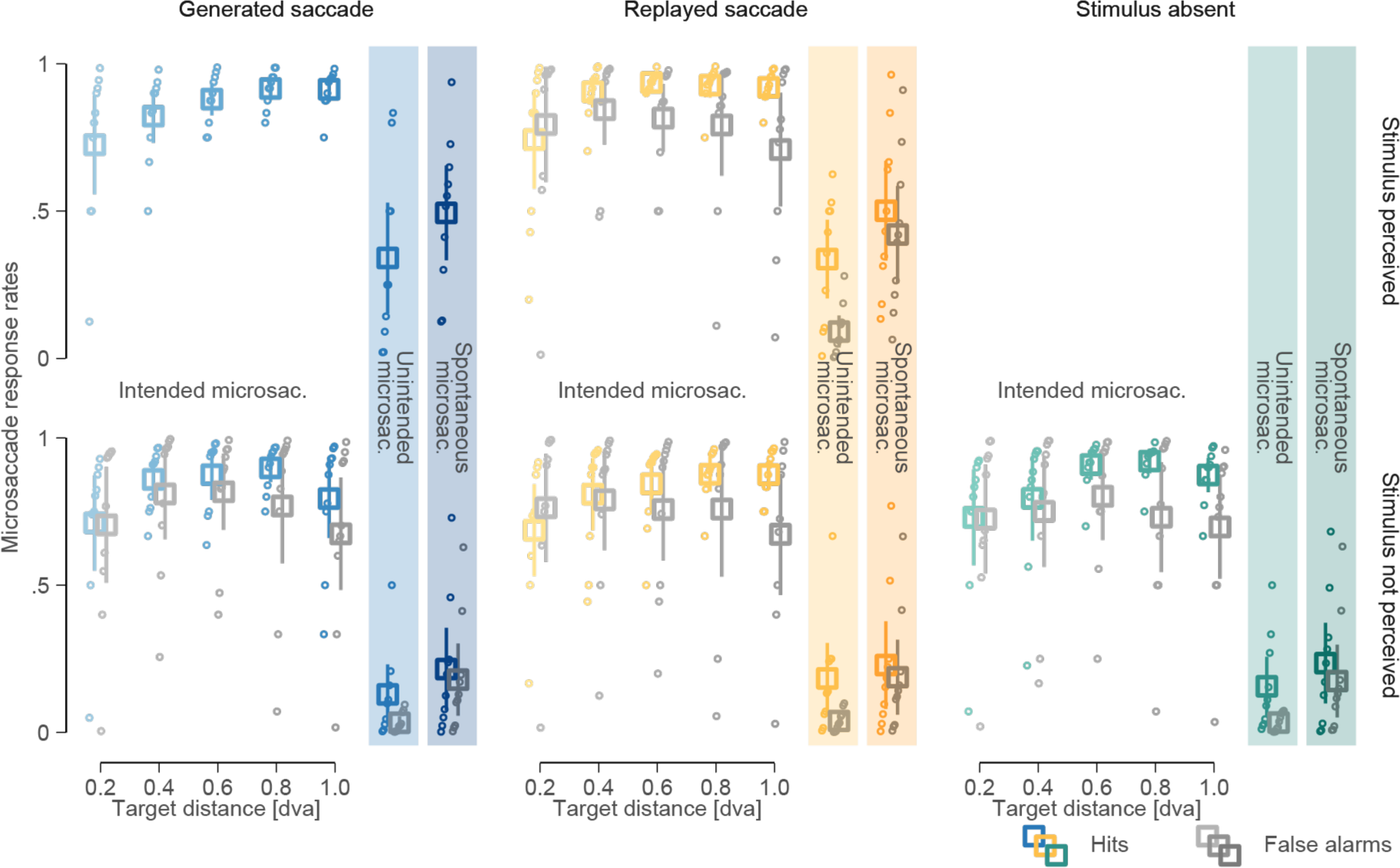
High rates for intended low rates for unintended and spontaneous microsaccades. Comparison of hit and false alarm rates for intended (**Exp. 1**), unintended (**Exp. 1**), as well as spontaneous microsaccades (**Exp. 2**). The data is split into different panels according to stimulus presentation condition (generated microsaccades in blue, replayed saccades in yellow, and stimulus-absent condition trials in green hues) and according to perceptual reports (stimulus perceived in upper, stimulus not perceived in lower panels). Data for intended microsaccades is additionally presented over five different target distances (ranging from 0.2 to 1 dva).

Finally, we examined if target distance affected detection of generated and replayed intended microsaccades in different stimulus conditions and depending on perceptual reports. Our three-way rmANOVA revealed a significant main effect of target distance (*F* (1.6,12.6) = 0.39, *p* = 0.047; test results are reported after Huynh-Feldt correction for violation of sphericity), and insignificant main effects of stimulus condition (*F* (1,8) = 1.29, *p* > 0.250) and perceptual report (*F* (1,8) = 2.52, *p* = 0.151). Interactions were all insignificant (all *p*s > 0.062). Post-hoc t-tests revealed significant differences between hit rates in trials with a target distance of 0.2 dva and all four remaining target distances (all Bonferroni-corrected *p* <= 0.011). No other comparison reached significance (all remaining Bonferroni-corrected *p* > 0.111). A two-way rmANOVA for false alarms in all stimulus conditions but only those trials for which participants reported no stimulus revealed no significant effects (all *p*s > 0.250). Together, these results indicate that, while hit rates were positively affected by saccade amplitude—with hit rates being the lowest when saccades are the smallest—false alarm rates stayed constant over the target distances.

